# Protein Fitness Prediction is Impacted by the Interplay of Language Models, Ensemble Learning, and Sampling Methods

**DOI:** 10.1101/2023.02.09.527362

**Authors:** Mehrsa Mardikoraem, Daniel Woldring

## Abstract

Advances in machine learning (ML) and the availability of protein sequences via high-throughput sequencing techniques have transformed our ability to design novel diagnostic and therapeutic proteins. ML allows protein engineers to capture complex trends hidden within protein sequences that would otherwise be difficult to identify in the context of the immense and rugged protein fitness landscape. Despite this potential, there persists a need for guidance during the training and evaluation of ML methods over sequencing data. Two key challenges for training discriminative models and evaluating their performance include handling severely imbalanced datasets (e.g., few high-fitness proteins among an abundance of non-functional proteins) and selecting appropriate protein sequence representations. Here, we present a framework for applying ML over assay-labeled datasets to elucidate the capacity of sampling methods and protein representations to improve model performance in two different datasets with binding affinity and thermal stability prediction tasks. For protein sequence representations, we incorporate two widely used methods (One-Hot encoding, physiochemical encoding) and two language-based methods (next-token prediction, UniRep; masked-token prediction, ESM). Elaboration on performance is provided over protein fitness, length, data size, and sampling methods. In addition, an ensemble of representation methods is generated to discover the contribution of distinct representations to the final prediction score. Within the context of these datasets, the synthetic minority oversampling technique (SMOTE) outperformed undersampling while encoding sequences with One-Hot, UniRep, and ESM representations. In addition, ensemble learning increased the predictive performance of the affinity-based dataset by 4% compared to the best single encoding candidate (F1-score = 97%), while ESM alone was rigorous enough in stability prediction (F1-score = 92%).

## 1. Introduction

Proteins are biological machines involved in almost all biological processes [1–4]. These molecules are made of amino acids that fold into 3-dimensional structures and perform life-sustaining biological functions [5]. Protein engineering practices aim to modify proteins to redirect what has already evolved in nature and address the industrial and medical needs of modern society [6,7]. This has been a challenging task due to the astronomical number of possible mutations and the complex sequence-function relationship of the proteins (i.e., fitness landscape) [8]. Therefore, various experimental and computational techniques have been developed to overcome the challenge of finding high-fitness proteins among mostly non-functional mutants. Recently, machine learning (ML) has shown promise as a tool to supplement already established techniques, such as rational design and directed evolution [9–12]. Unlike directed evolution, ML models can learn from non-functional mutants instead of simply discarding them during enrichment for functional clones. ML-assisted protein engineering, therefore, has potential as a time-efficient and cost-effective approach to searching for desired protein functionality. This provides a unique opportunity to create smart protein libraries, elevate and accelerate directed evolution strategies, and enhance the probability of finding unexplored high-fitness variants in the protein fitness landscape [13–15]. Machine learning methods have attained a high success rate in predicting essential protein properties including secondary structure, solubility, binding affinity, flexibility, and specificity [16–20]. Despite these recent milestones, generalizability and robustness of models will require further explorations in different protein fitness prediction tasks and training details.

Solving these challenges will require us to view proteins from a new perspective that supplements our biochemical knowledge with lessons from written languages. Recent advances in ML and artificial intelligence have applied natural language processing (NLP) methods to identify context-specific patterns from written or spoken text. NLP tasks learn how words function grammatically (syntax) and how they deliver meaning within themselves and in surrounding words (semantics) [21,22]. This has given rise to virtual assistants with voice recognition and sentiment analysis of text from diverse languages [23,24]. Similarly, protein engineering can leverage these NLP tools – treating a string of amino acids as if they were letters on a page – to understand the language of proteins, providing a promising route to capture nuances (e.g., epistatic relationships, functional motifs) in complex sequence-function mappings [25,26]. The rapid expansion of publicly available protein sequence data (e.g., Uniprot [27], SRA [28]) further supports the use of big data and language models in the domain of protein engineering[29]. Self-supervised language models learn the context of the provided text by reconstructing the masked tokens/linguistic units of the text string using the unmasked parts. For the context of protein engineering, pre-trained protein language models – carrying valuable information about the epistasis/interaction of amino acids – can be applied to downstream tasks by extracting the optimized weight functions as a fixed-size vector (embedding) [25,30,31]. Among early embedding developments, Alley et al. introduced UniRep, a deep learning model that was trained on 24 million unique protein sequences to perform the next amino acid prediction tasks for extracting information about the global fitness landscape of proteins. Rives et al. trained ESM, a language model for masked amino acid prediction tasks, on over 250 million protein sequences [32]. The learned representations – including UniRep [33], ESM [32], TAPE [34], and ProteinBERT [35] *-* have generated promising results in diverse areas such as predicting protein fitness, protein localization, protein-protein interaction, and disease risk of mutations in terms of improved prediction scores, increased generalizability, and mediated data requirements [36–40]. Using embeddings for sequence representations (transfer learning) enables knowledge transfer between protein domains and future prediction tasks by further optimizing the already-learned weights. For example, Min et al. obtained a 20% increase in the F1-score (the harmonic mean of precision and recall) for a heat shock protein identification task when training their NLP-based model, DeepHSP [41], on top of pre-trained representations.

In this study, we perform protein sequence fitness prediction with ML techniques to demonstrate how model performance varies given the choice of protein representation, protein size, and the biological attribute (e.g., binding affinity and thermal stability) to be predicted. This work provides actionable insights for effectively building discriminative models and improving their prediction scores via sampling techniques and ensemble learning. As efficient use of embedding methods on experimental datasets is in its infancy, rigorous studies are needed to gain new insights into the performance of the pre-trained models given various training conditions and distinct biological function predictions. To this end, we two large datasets that were representative of common protein engineering tasks. First, we leverage a highly imbalanced dataset (93% non-functional; Table 1), consisting of our previously described affinity-evolved affibody sequences [42] to explore NLP-driven practices. We then expand our analysis to include thousands of protein sequences labeled with their experimentally measured stabilities (melting temperatures, Tm) obtained from the Novozymes Enzyme Stability Prediction (NESP) dataset [43]. With the two datasets having unique attributes, we were well positioned to address multiple questions: i) How do different representation methods perform in predicting distinct fitness attributes such as stability or affinity? ii) How do sampling methods perform in the imbalanced protein dataset? iii) Is ensemble learning over different protein representations helpful in boosting the performance of discriminative models? By addressing these challenges, we also gain direct insights for model interpretation and reveal the features that are most important for discriminating between fit and non-fit sequences (Figure S1, Figure S3-4). We discovered that oversampling generally outperformed the undersampling techniques (Figure 4). In addition, ensemble over representations greatly improved the predictive performance in the affibody data (Figure 5). For protein representations, UniRep and One-Hot outperformed other methods in the affibody (affinity) dataset while ESM achieved the best score in stability prediction in NESP (Figure S6). Finally, it was observed that the performance of various protein representation methods is strongly impacted by protein sequence length (Figure 7).

**Table 1.** Dataset attributes and prediction tasks

| Dataset | Task | Fitness | Model | Attributes |
| --- | --- | --- | --- | --- |
| **Affibody** | Classification | Binding Affinity | Logistic Regression | 82,663 non-binders<br>6,077 binders |
| **NESP** | Classification | Stability | Logistic Regression | 3,743 high-stability<br>1,311 low-stability |
| **NESP** | Regression | Stability | Random Forest Regressor | 18,190 total |

## 2. Materials and Methods

### 2-1 Obtaining Experimentally Labeled Sequence Data

Two different datasets with varying data characteristics were explored. The first is our experimental data of affibody sequences that previously were iteratively evolved for binding affinity and specificity against a panel of diverse targets [44]. The second collection of labeled protein sequences was obtained from the recently released Kaggle dataset wherein numerous proteins (n= 18,190) of various lengths are labeled according to their thermal stability (Tm). The NESP dataset was filtered to only include sequences characterized at pH=7. For the affibody dataset, raw sequence data were cleaned by removing any sequences that contained stop codons or invalid characters. Afterwards, the frequency of each unique sequence in the experimental steps was tabulated. Infrequent sequences appearing fewer than ten and four times (within magnetic activated cell sorting (MACS) and fluorescent activated cell sorting (FACS), respectively) were treated as background and removed from the analysis. Note that more stringent frequency removal for MACS was mainly due to the experiment type and higher probability to introduce noise in the dataset. After removing the background, sequences from MACS and FACS were combined to form the final high-fitness population of binders. The non-binding population included the initial affibody sequence pool which did not appear in the enriched population of binder sequences. The Initial affibody sequences that were within one hamming distance (i.e., a single amino acid mutation) of any enriched sequence were removed as well to account for potential errors encountered during deep sequencing. All affibody sequences were exactly 58 amino acids in length with mutations present at up to 17 of these positions.

### 2-2 Obtaining the Sequence Representations

We obtained four different numerical representations for our sequence data: One-Hot and physiochemical encoding, UniRep, and ESM embeddings. One-Hot encoding refers to building a matrix (amino acids × protein length) and filling it with one when there is a specific amino acid in the given position, filling the rest with zeros. For physiochemical encoding, we used the modlamp [45] package in python which is used for extracting the physical features from protein sequences. There were two types of features represented in modlamp package (global and peptide descriptors). All the global (e.g., sequence length, molecular weight, aliphatic index, etc.) and local physiochemical features based on Eisenberg scale were extracted for this analysis (twenty in total).

Embedding refers continuous representation of the protein sequence in a fixed-size vector, and it should contain meaningful information about proteins [46]. For example, in the embedding visualization of amino acids in low dimensions for both UniRep and ESM, similar amino acids (in terms of size, charge, hydrophobicity, etc.) were close to each other. For UniRep representation, we used the 1900 dimension and mean representation over layers. We used Jax_UniRep since for obtaining the UniRep embeddings, https://github.com/ElArkk/jax-unirep. UniRep uses mLSTM structure for performing next-token prediction, and it was trained on 24 million sequences in the Uniref50 dataset with 18M parameters. For ESM, we chose ESM2 [47] with 1280 vector dimensions and 650M parameters and means over layer representations. GitHub for ESM is https://github.com/facebookresearch/esm.

### 2-3 Sampling & Splitting

Sampling refers to choosing a random subset of data to represent the underlying population. Three different sampling methods were tested for our severely imbalanced affibody dataset: undersampling, random oversampling, and synthetic minority oversampling technique (SMOTE) [48]. Due to the sparse and rugged nature of the protein fitness landscape, it is common for experimental data obtained in the protein domain to be highly imbalanced. One practical approach for resolving the imbalanced dataset issue is using sampling techniques when training the dataset. Oversampling is randomly repeating the minority class examples; thus, it could be prone to overfitting in comparison to undersampling. However, undersampling may discard the useful information especially in severely imbalanced datasets as it is removing many samples from the majority class. SMOTE is a more recent addition to sampling methods, and it is oversampling the minor population by synthetically generating more instances that are highly similar to the minority class. While SMOTE has shown promising results in increasing the prediction performance for various imbalanced datasets [49–51], there are also studies indicating undersampling superior performance compared to oversampling methods [52,53]. As a result, we examined the performance of all three sampling techniques to validate which sampling method performs well within our wet-lab protein dataset over different encoding methods.

For splitting the datapoints within the test set in an imbalanced dataset, sampling equally from each class may lead to an overestimation of the model performance[54]. As a result, we made sure that the test set distribution follows the initial data distribution (93% naïve vs. 7% Enriched). The classification performance was examined with F1-score. Note that hyperparameter optimization was implemented when necessary with OPTUNA [55] and the objective was set to maximize the F1-score in validation set.

### 2-4 Algorithm Selection & Training Details

For classification, logistic regression (LR) was chosen and L2 penalization (Ridge) was used to reduce the likelihood of overfitting. We reasoned that a simple logistic regression enables a fair comparison between cases. One regression task was also implemented over the NESP dataset with random forest regressor (RFR). We used regression to observe how models perform with increasing the prediction challenge, from binary prediction to actual label prediction. The rationale for using RFR was that linear regression model was not viable to meet the prediction task complexity. For a fair comparison between protein encoding performances in regression, the RFR hyperparameters, max number of estimators and max_depth, were optimized with OPTUNA.

### 2-5 Ensemble Learning

To improve the predictive performance of protein encoding predictions, we developed a framework that combines various encoding methods. We experimented with two approaches: **concatenation and voting**. In concatenation, the encodings were combined by adding them together and used the resulting representation as input for our predictive model. In voting, separate predictive models for each encoding method were trained. The final prediction is then calculated with the majority-voted label over a fixed test set.

### 2-6 Metrics & Statistical Analysis

The metric used for analyzing classification performance is the F1-score which quantifies the prediction power even in imbalanced datasets as it takes both precision and recall into account. For regression, we used mean squared error (MSE) and R^2^ to indicate how the models perform. MSE is the mean of the square of differences between the actual labels and the predicted values in the test set while R^2^ represents the variation explained by the independent variables. For sensitivity analysis, experiments were implemented with multiple random seeds (20 in affibody and 30 in NESP dataset) and p-value has been calculated to analyze the null hypothesis. The null hypothesis assumes that the performances of the methods are similar and when rejected, we consider the methods to be statistically significant in their obtained output. The results for multiple seeds are shown with violin plots where the white dots represent the mean value.

The techniques outlined in this section are summarized in Figure 1.

**Figure 1.**
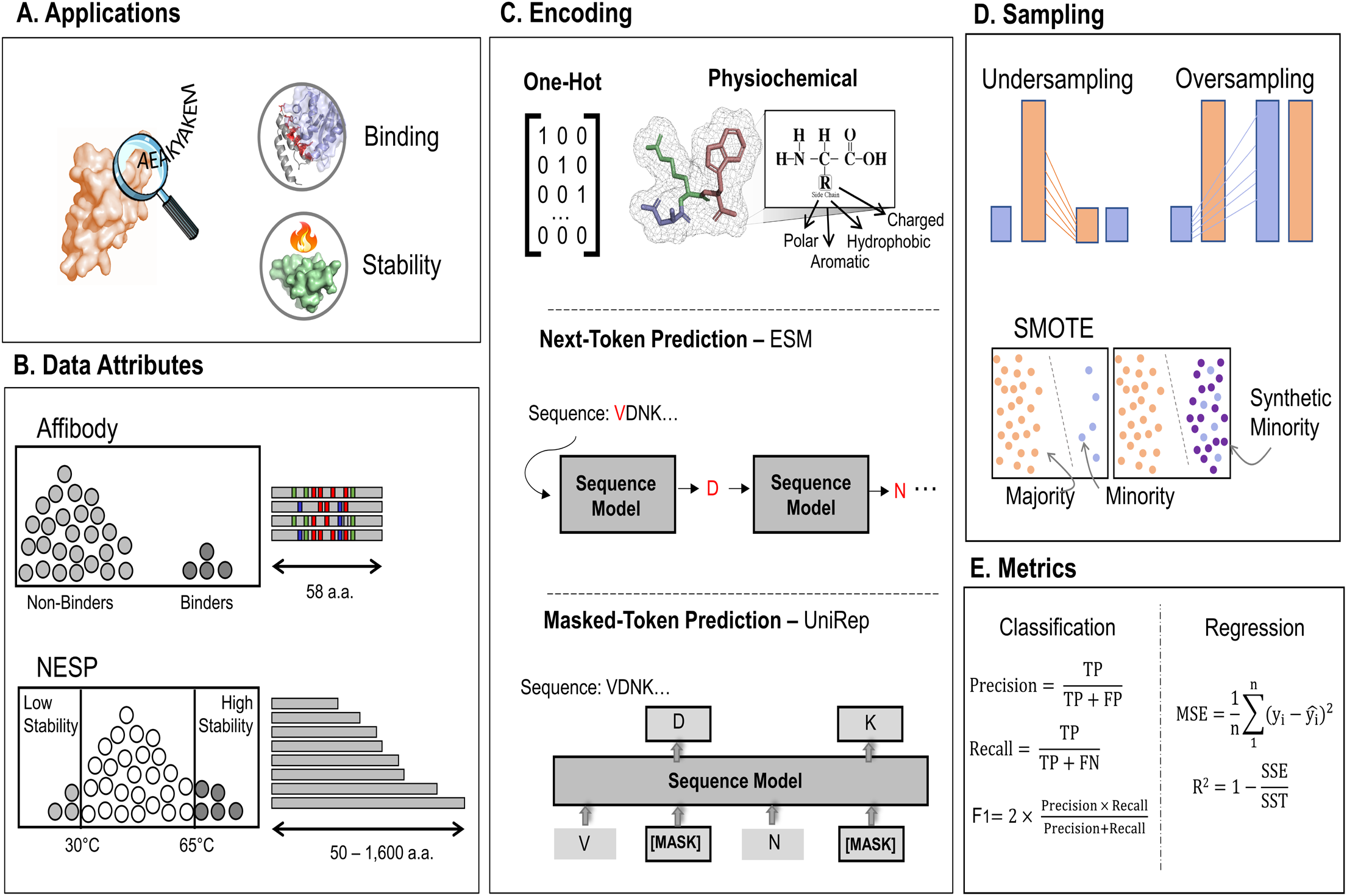
Overview of the implemented techniques, data attributes, and evaluation metrics. A. Illustrates the use of sequence-function mapping to identify protein sequence functionality (e.g., therapeutics, diagnostics, enzymatic function). B. Data attributes for the two datasets used in this study. The first dataset includes high-fitness protein binders among a pool of non-binder affibody sequences with up to 17 mutation sites. The other dataset includes a wide array of proteins with their associated melting point. C. One-hot encoding, physicochemical encoding, and pre-trained models were used to encode the protein sequences present in our datasets. All present protein amino acid information in a machine-readable format, but in different ways. One-hot encoding converts each amino acid to a binary vector of all 0s but 1 where it belongs to its position in the matrix. In physicochemical encoding, each amino acid is represented by its physiochemical characteristics, such as polarity, charge, size, etc. Pretrained models are trained over a large corpus of unlabeled data capturing the syntax and semantics of protein language via NLP-driven models, such as next-token prediction (e.g., UniRep) and masked token prediction (e.g., ESM). D. The sampling methods to be discovered in this literature are undersampling, oversampling, and synthetic minority oversampling techniques (SMOTE). E: The metrics used for evaluating the performance of prediction tasks (classification and regression) are defined.

## 3. Results

### 3.1. Sequence-Function Mapping Obtained from High-Throughput Selection Methods & Deep Sequencing Affibody Dataset

To investigate the impact of feature representation, ensemble learning, and sampling methods, several prediction tasks were leveraged. We performed a classification task on the obtained sequences to predict the scarce high affinity binder class among the pool of non-binder class in the affibody data. For NESP dataset, in the classification task, we simplified the data by choosing two classes of low- (Tm ≤35°C) and high-stability (Tm ≥60°C). In addittopm, regression was implemented to increase the prediction difficulty and to observe how protein encodings perform releatively. The models were tasked with predicting the stability (Tm) value and all the sequences with measured pH=7 were included. The details of obtained sequences after cleaning, and the type of prediction tasks are reported in Table1. **Note that the NESP results will be overviewed in section 3-5 and supplementary information**.

### 3.2. Physiochemical Feature Encoding, Interpretable yet Lower Predictive Capacity

The classification results in physiochemical encodings are shown in Figures 2 and 3. We ranked the leading features in discriminating non-binder and binder classes and listed the encoding method’s F1-score in different sampling methods. The physiochemical encoding performance was not among the lead encoding methods, yet it achieved a high F1-score with only 20 features. It also provided insights on how physiochemical features correlate with each other in the given data (Figure S1).

**Figure 2.**
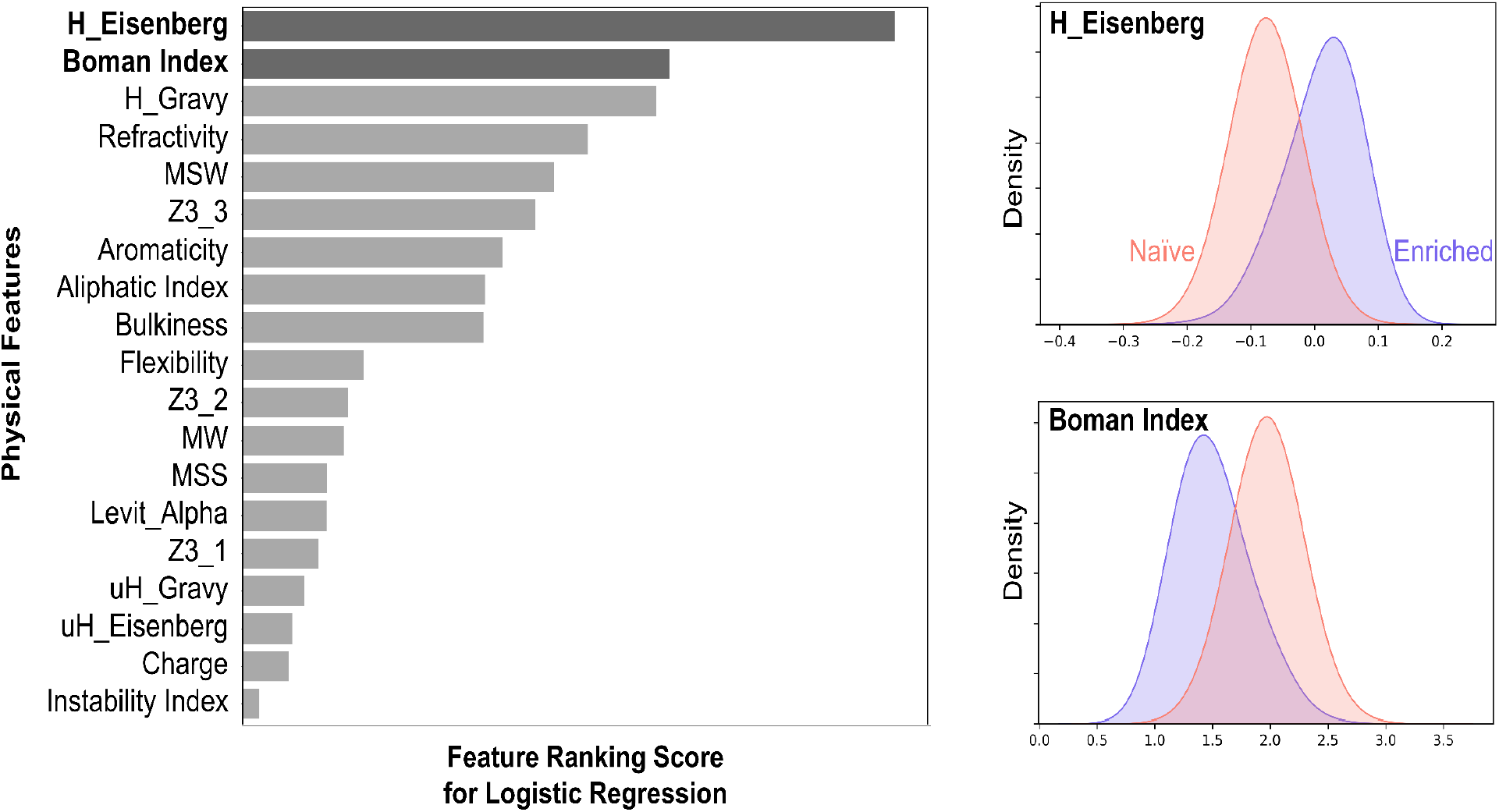
The lead physiochemical features in naïve and enriched class discriminations were H_Eisenberg, Boman Index, and H_Gravy. Gravy and Eisenberg capture hydrophobicity scales. Boman Index is a measure of the protein’s ability to interact with its environment based on the solubility of individual residues. The enriched proteins in our library have gone through negative screening and are specific to their target. Therefore, there is a shift to a lower Boman index for this population. Note that the plot is the result of oversampling, SMOTE, in the logistic regression task.

**Figure 3.**
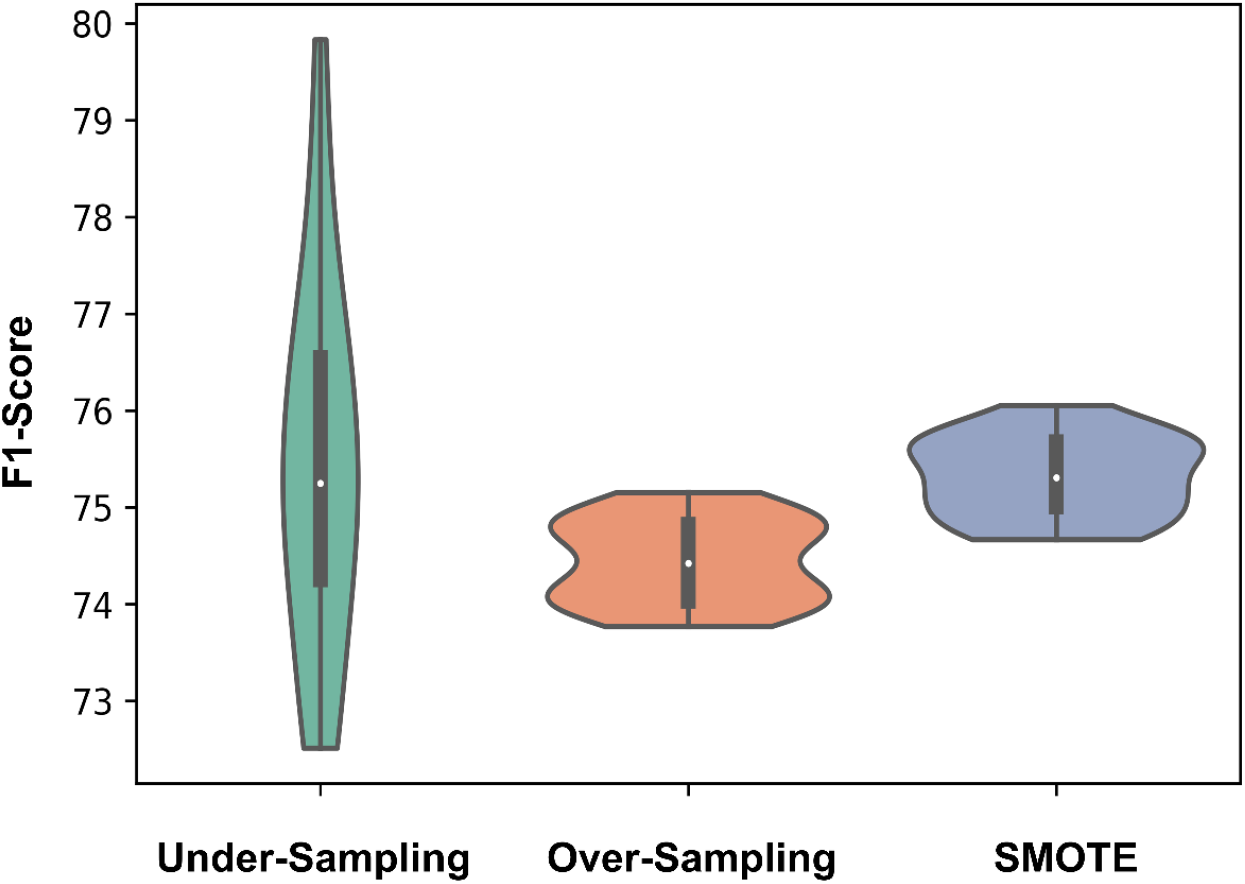
SMOTE with mean F1-score= 75.5% was found to be the most effective sampling method when encoding the affibody sequences with physiochemical features. A) Sensitivity analysis of the predictive performances with 20 random seeds. The results from the analysis indicate that undersampling might perform as powerfully as SMOTE since the p-value>0.05. However, incorporating all the p-values, SMOTE has shown a highly significant difference in F1-score compared to oversampling.

### 3.3. Comparison Over All the Encoding and Sampling Methods

Once the lead physical features for high-affinity binders were determined, we demonstrated the performance of different protein representations within our selected sampling techniques. The prediction performance indicates that each encoding method performed differently in predicting the fitness of proteins, and One-Hot and UniRep were the top performers. In addition, among the samplings, SMOTE boosted the F1-score in almost all cases. Figure 4 exhibits the F1-score distributions within 20 different random seeds. A complete T-Test analysis is reported in Table S1-3.

**Figure 4.**
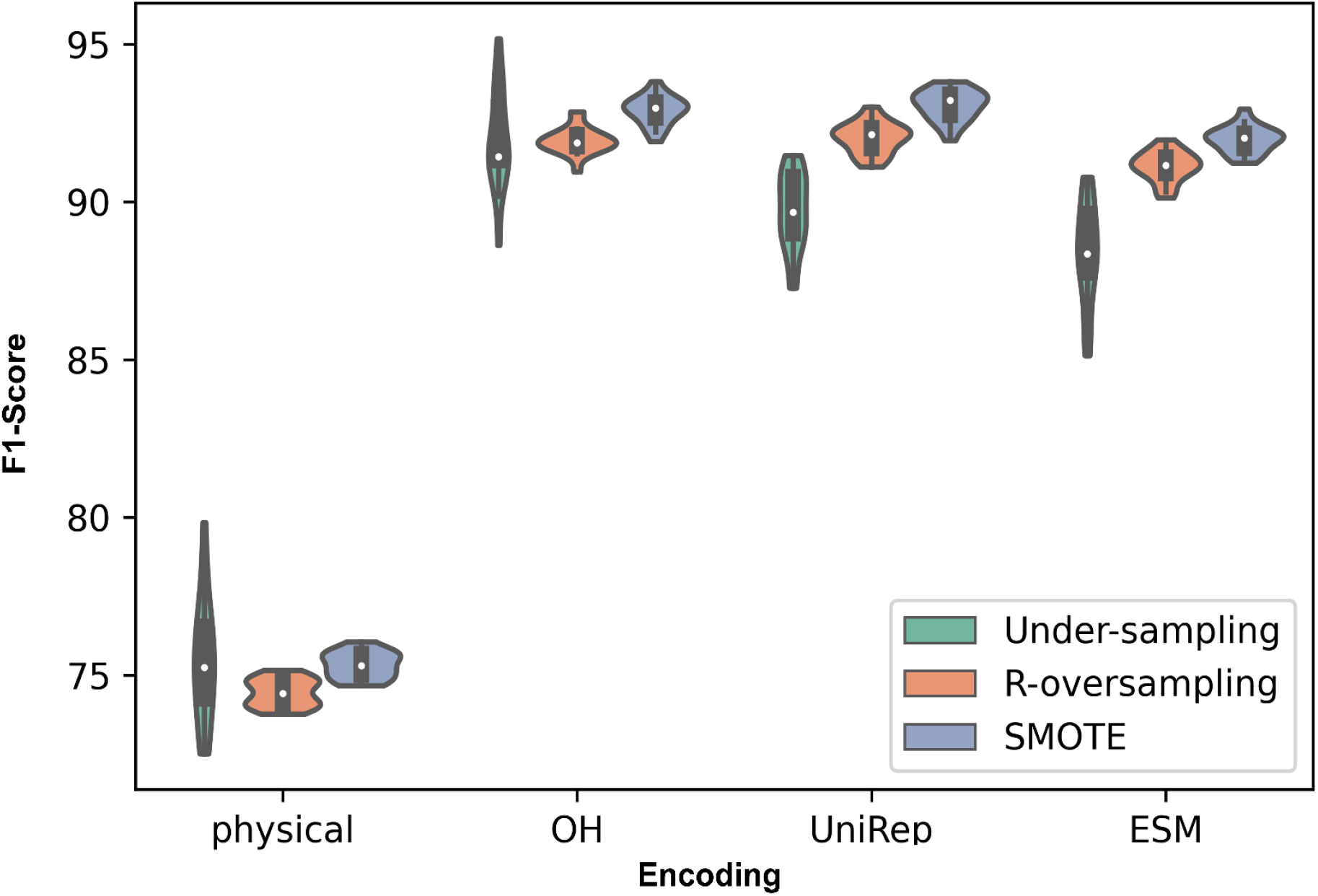
Performance analysis of encoding methods highlights the shortcomings of physical features and strength of the SMOTE sampling method. Protein sequences encoded using physical features, one-hot, UniRep, and ESM were used to perform classification tasks among the affibody dataset. Within each encoding method, under-sampling, random over-sampling, and SMOTE sampling methods were evaluated for each encoding. The resulting F1 scores over 20 random seeds are shown here as violin plots. Detailed statistical analysis is provided in Table S1.

### 3.4. Increased Generalizability & Predictive Performance via Ensemble Learning

Due to the varying performances of the protein encodings, we postulated that ensemble learning increases the models’ predictive performance. As oversampling performed better than undersampling in three out of four encoding methods, we exclusively analyzed the ensemble learning for the two oversampling types (i.e., R-oversampling, and SMOTE). The physicochemical encoding for this analysis was discarded as its performance was not as potent as the other encodings.

Figure 5 represents ensemble technique, voting, remarkably enhanced the performance with respect to all the methods with a mean F1-score=97% over the 20 random seeds. This represented voting method enhanced the prediction score by combining the predictions of multiple models based on single encodings. We concluded that as different encodings might capture distance and relationship of the datapoints differently, combining their predictions boosted the final model performance. The encoding methods used for voting technique in the dataset are visualized in Figure 6 in a UMAP (Uniform Manifold Approximation and Projection) [56] plot

**Figure 5.**
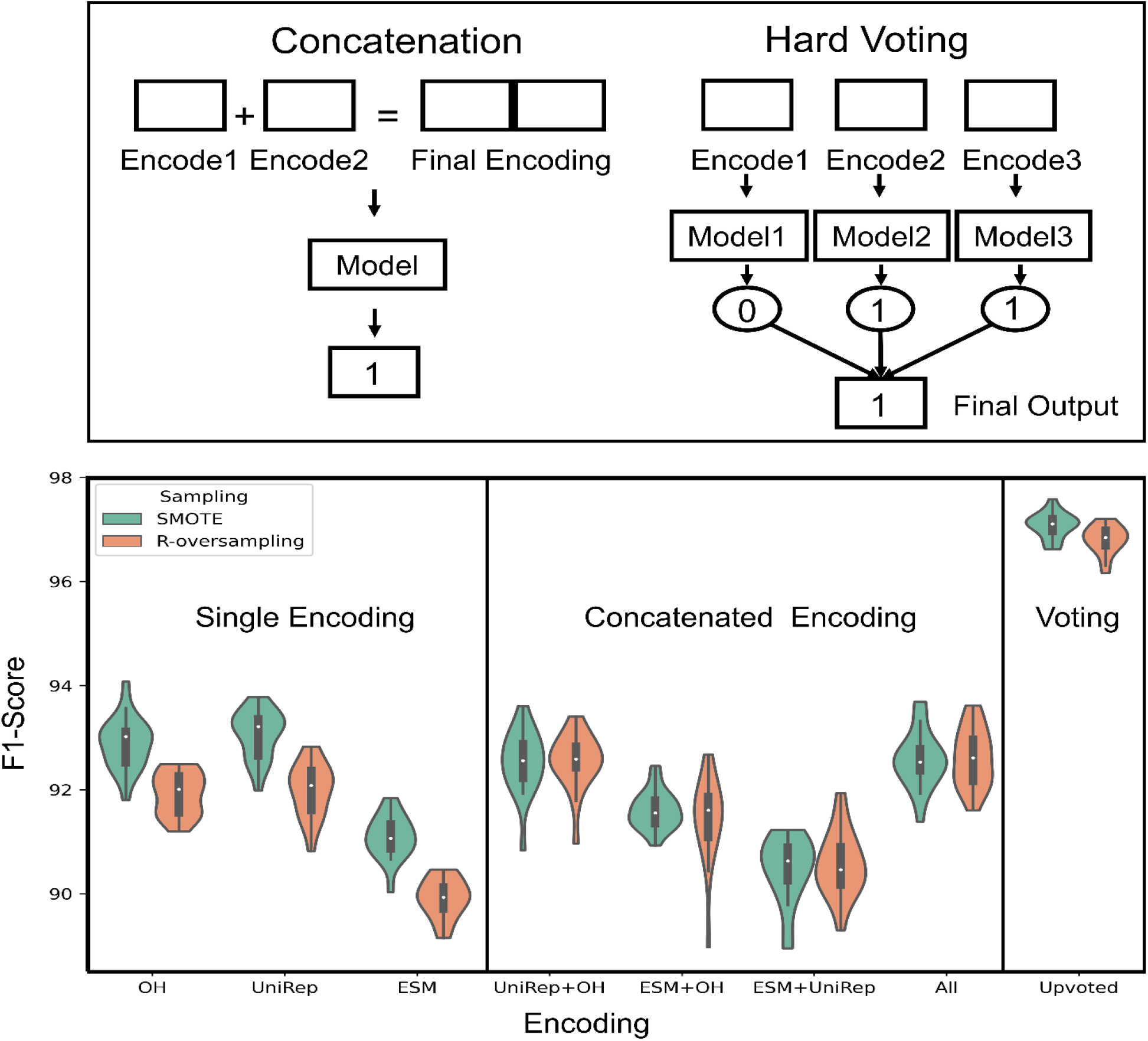
Voting Substantially Improved the predictive performance in All Random Initializations over Different Encoding Methods. The plot above has three regions from left respectively; it includes single encoding methods, concatenation of encodings, and voting of predictions. The vote was performed such that each encoding went through a predictive model over the same dataset. Then, the final prediction is obtained by majority voting. It is insightful how voting increases the models’ robustness and generalizability. The concatenation performed similarly or worse than the best model in single encodings. For the statistical analysis, please refer to the supplementary. The best model among all predictions was Upvote with SMOTE sampling, Mean-F1-score=97%. Refer to the supplementary for a summary of statistics (Table S2).

**Figure 6:**
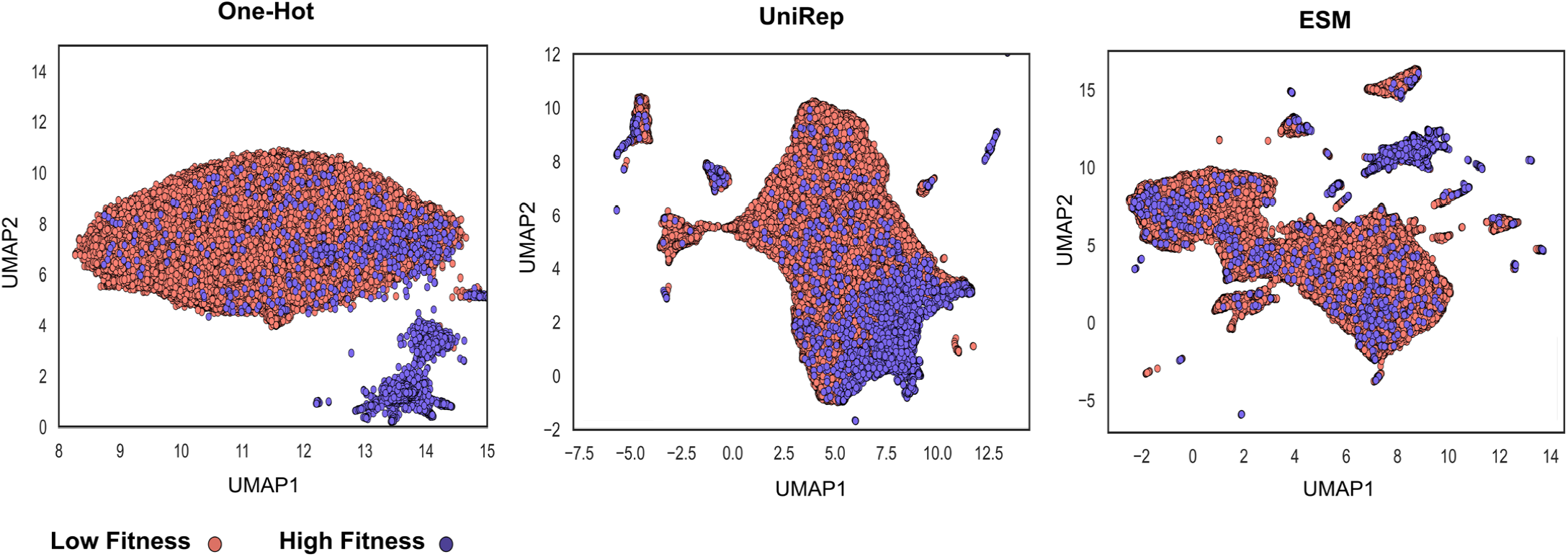
Different protein encodings potentially capture distinct functional aspects of the proteins. A 2D visualization of the encoding techniques which resulted in improved prediction in the voting method in UMAP. This method is a dimensionality reduction technique such as PCA with unique advantages such as preserving the local structure of the data and capturing non-linear relationships between data points. In observing the sequence-function relationship in proteins, one can conclude that each protein sequence representation/encoding has the potential to capture different aspects of the fitness to be predicted.

Note that while F1-score is used as an overall measure of model’s performance, individual analysis of confusion matrix values of a classification algorithm affords clarity on how the model performs predicting the labels for classes (Figure S2).

### 3.5. How Protein Encodings Perform Considering the Data Attributes

The hypotheses were tested over affibody datasets that had notable attributes such as severe imbalance, multiple mutation sites, affinity and specificity enrichment, and small molecular protein length. The obtained results indicated voting and oversampling were highly effective methods to boost the fitness prediction performance. However, individual protein-encoding performance comparisons need more convincing explanation and thorough exploration. Specifically, we wondered why ESM underperformed one-hot and UniRep despite more powerful setup in pretraining and a being showcased in studies for high prediction potential [57]. While the performance could be due to the datatype (e.g., small protein, complex fitness, etc.), we decided to further analyze the encoding prediction scores in a completely different dataset and bring insights on embedding performances in various conditions (e.g., data size in training, protein length, prediction task difficulty). The curated data contains 18,190 sequences with varying lengths and provides melting points which indicate the protein stability. Figure 7 is the performance comparison in stability prediction of embeddings, their concatenation, and voting using different data sizes. Despite down performing in the affibody affinity data, ESM performed best for stability prediction when including proteins with max length=500 (Figure S6).

**Figure 7.**
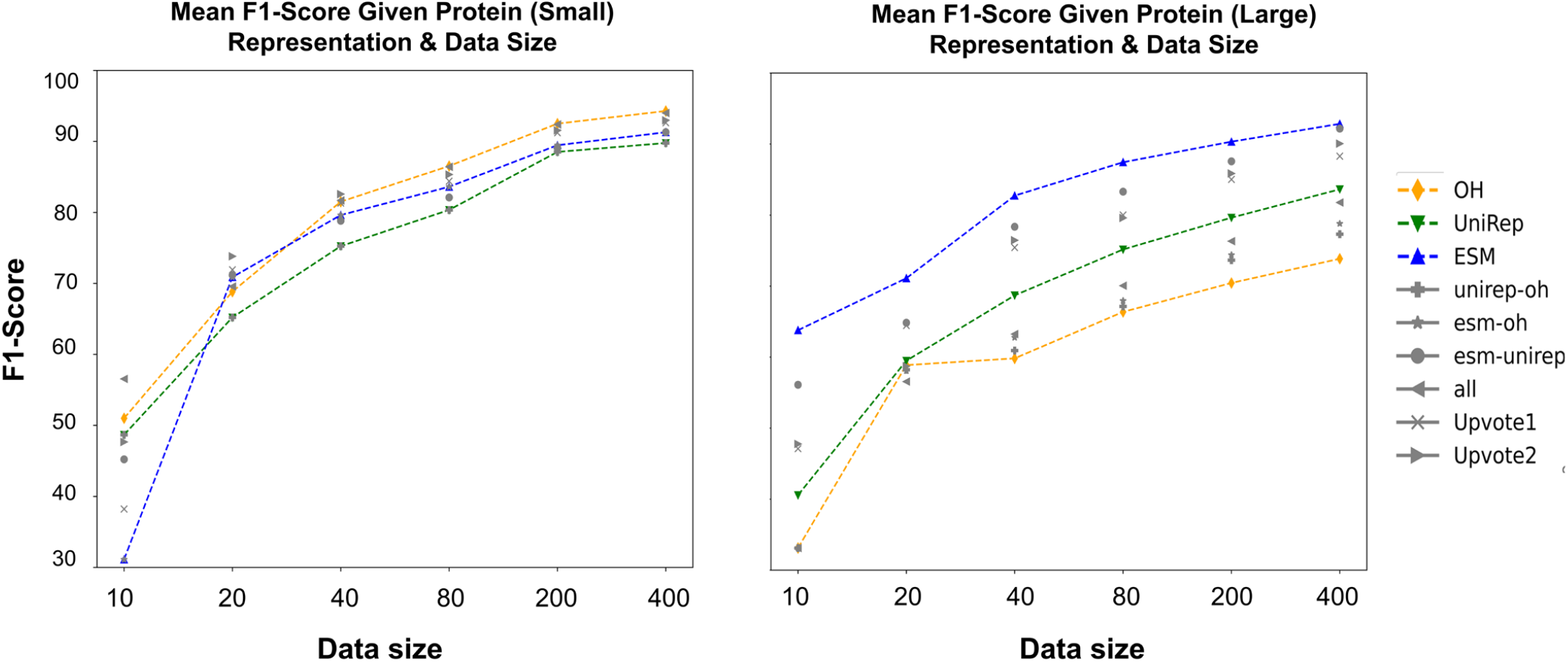
Protein representation performances over different data sizes for small proteins (length<=120) vs. large(400<=length<=1500). The obtained results are largely different with respect to the protein size. **Highlights: For small proteins**, upon comparing the violin plots and statistical test results, protein sequence encoding methods were performed distinctively with respect to the initial dataset (protein max length=500). One-Hot encoding had a more significant contribution in boosting the classification metrics for small proteins. As an example, when N=400, both One-Hot and All-Encoding concatenation with a mean F1-score of 94% outperformed the other encoding methods. **One-Hot tends to be problematic for large proteins** as it results in a highly sparse encoding vector. This has been shown in this plot when One-Hot encoding performance is not satisfactory enough in comparison with ESM and UniRep. When N=400, Based on both the violin plots and Welch t-test, either ESM or ESM_UniRep with 92% mean F1-score achieved the highest performance. One-Hot with *a* 73% mean F1-score was the lowest score among all the encodings. Refer to the supplementary information for all one-by-one comparisons of the statistics and classification scores.

The last analysis is a regression task for predicting the melting point value. We wondered how different encoding methods perform if we use all the data and increase the prediction challenge (Tm prediction rather than stability class prediction). MSE and R^2^ are shown in predicting the Tm values of a dataset of 18,190 sequences with 0.3 test size. There was a significant difference in the performance of encoding methods which was not the case in classification task. ESM was the best encoding method in predicting stability (R^2^ score= 0.65). Note that we used all the data and did not use sampling methods for training. Therefore, sensitivity analysis was not critical to perform for performance estimations. The regression metrics are reported in Table2.

**Table2.** Regression metrics for encoding methods in validation & test set.

| Encoding | Validation |  | Test |  |
| --- | --- | --- | --- | --- |
|  | R<sup>2</sup> | MSE | R<sup>2</sup> | MSE |
| One-Hot | 0.21 | 141 | 0.24 | 130 |
| UniRep | 0.49 | 108 | 0.4 | 102 |
| ESM | 0.65 | 63 | 0.65 | 60 |

## 4. Discussion

In this study, we shed light on two key challenges of applying discriminative models over amino acid sequence data for protein engineering applications: 1) handling imbalanced data and 2) choosing an appropriate protein representation (i.e., encoding). Assay-labeled sequence data in this domain is often severely imbalanced (due to the rugged and sparse nature of the protein fitness landscape) and requires careful consideration in data sampling, splitting, and choice in data representation for model training. To capture this common occurrence of imbalanced data, we trained discriminative ML models over our cytometry-sorted deep-sequenced small protein (affibody) data to distinguish between functional sequences (n=6,077) among a large collection of non-functional protein sequences (n=82,663). We then explored the impact of encoding protein sequences using two simplistic approaches (One-Hot encoding, physiochemical encoding) and two language-based methods (UniRep, ESM). We hypothesized that, as each protein representation may capture distinct information, combining representations via embedding concatenation and ensemble learning increases overall performance and generalizability.

To address the issue of imbalanced data, we implemented multiple sampling techniques – undersampling, random oversampling, and SMOTE – and compared performances via F1 scores. Our results indicate that implementing oversampling techniques over imbalanced datasets improves predictive performance relative to undersampling or the exclusion of sampling methods. Among the sequence representation methods, embeddings are the answer to improved fitness prediction and data requirements. However, it is essential to consider the choice of protein representation, its benefits, and its drawbacks. For example, the choice of fitness to be predicted (e.g., thermal stability, binding affinity, target specificity) and the language model pretraining procedure affect the model’s predictive performance and need further discussion. Therefore, we analyzed an additional dataset (i.e., the NESP dataset which included a variety of protein sequences with their Tm) to discuss the effect of protein representations over the variables such as protein length, protein fitness, and prediction type (i.e., classification vs. regression). For ensemble learning, we used majority voting to combine the prediction of each representation over the same ML model which significantly improved the F1-score in Affibody dataset.

As only a very small fraction of protein sequences is experimentally annotated with properties, the primary goal of embeddings is to distill valuable information from unlabeled data and use them for property/fitness prediction. Previous reports have observed that there are sequence motifs, conserved regions, and evolutionary information in the protein databases that can be learned by language models [33,34,58]. This has been tested with different NLP techniques, varying model parameters, and clustering sizes for databases used and resulted in a wide array of language-based protein representations [30,57,59,60]. These promising embeddings (e.g., ESM, UniRep) have been evaluated in many studies and have improved the fitness prediction scores and alleviated the assay-labeled data requirements [59,61]. However, there are also studies that report minor improvements to predictions by using solely embedding methods. In some cases, prediction scores were improved by simpler representations such as one-hot or physiochemical encoding [62,63]. Similarly, Rao et al. pointed out a different performance of embeddings in TAPE [34] with 38 million parameters based on different protein engineering tasks. Their model performed outstandingly in fluorescence and stability prediction while it did not perform as well as hand-engineered features in contact prediction.

The current capabilities and limitations of language models motivate the need for optimizing the pretraining task and improving the methodology for supervising the pre-trained models. Consider ESM2, one of the largest language models used for protein sequences which has shown significant improvement in protein structure prediction compared to previous embeddings. In our study, protein representations obtained via ESM2 significantly outperformed UniRep or One-Hot in stability prediction. However, in the context of predicting binding functionality among small protein affibody variants, its performance was exceeded by UniRep and One-Hot (Figure 4). This motivates looking into what knowledge is transferred by pretraining models and how useful they are for specific fitness predictions, with or without further supervision. Here we covered the core challenges and considerations in supervising the models in fitness prediction, yet additional downstream analysis and posing insightful questions will give us more understanding and directions in discriminating the protein sequences based on their fitness. In order to improve the pretraining step, we might adopt techniques such as adjusting the masking rate [64], adding biological priors [60,65], increasing the model parameters [57], and building specialized language models for the desired fitness [66], given the growing data availability and computational resources. Additional studies are required for improved downstream fitness predictions, such as fine-tuning with a reduced chance of overfitting [67], incorporating the effect of post-translational modifications, and characterizing the performance of embeddings in different data setups [68] with varying protein types and finesses for supporting the development of novel proteins in diagnostics and therapeutics.

## 5. Conclusions

We integrated machine learning and protein engineering knowledge to identify high-fitness protein sequences. We quantified model performance while varying the choice of feature representation, ensemble learning, and sampling methods. Analysis across a broad range of protein chain lengths revealed the ESM language model to be most beneficial for encoding large protein sequences (Figure 7). Yet, in the context of small protein sequences, comparable performance was observed between one-hot encoding and the language models (ESM and UniRep). In our analysis, oversampling proved to be an effective technique to improve performance when dealing with severely imbalanced datasets (Figure 4). Finally, ensemble learning was a promising method for boosting the prediction scores when using unique, competitive encoding methods (Figure 5).

## Data and Code Availability

The NESP data is available at https://www.kaggle.com/competitions/novozymes-enzyme-stability-prediction/data. The Affibody dataset is available upon request. The source code for this project can be found On the GitHub repository: https://github.com/WoldringLabMSU/Sequence_Fitness_Prediction.

## Acknowledgments

We would like to thank the department of chemical engineering and material science at Michigan State University for funding this project. For the support in computational resources, we thank MSU’s High Performance Computing Center (HPCC). Finally, we would like to express our gratitude to Alex Golinski for his invaluable comments and insights.

## Supplementary Materials

The following supporting information can be downloaded at the provided link.

Figure S1: Physicochemical feature correlation plot affibody dataset.

Table S1: Figure 4 t-test results.

Table S2: Figure 5 t-test results.

Table S3: T-test results for R-Oversampling vs. SMOTE.

Figure S2: Violin plot-based confusion matrix for Figure 5 results.

Table S4: T-test for Figure S2.

Figure S3: Physicochemical feature correlation plot NESP dataset.

Figure S4: Physicochemical feature ranking for NESP dataset.

Figure S5: Physical Feature representation while using maximum N=1000, performed poorly and have not got selected for the main figure.

Figure S6: Mean F1-score comparison between protein representations including proteins with max length=500.

