## Supplementary Information for "Protein Fitness Prediction is Impacted by the Interplay of Language Models, Ensemble Learning, and Sampling Methods"

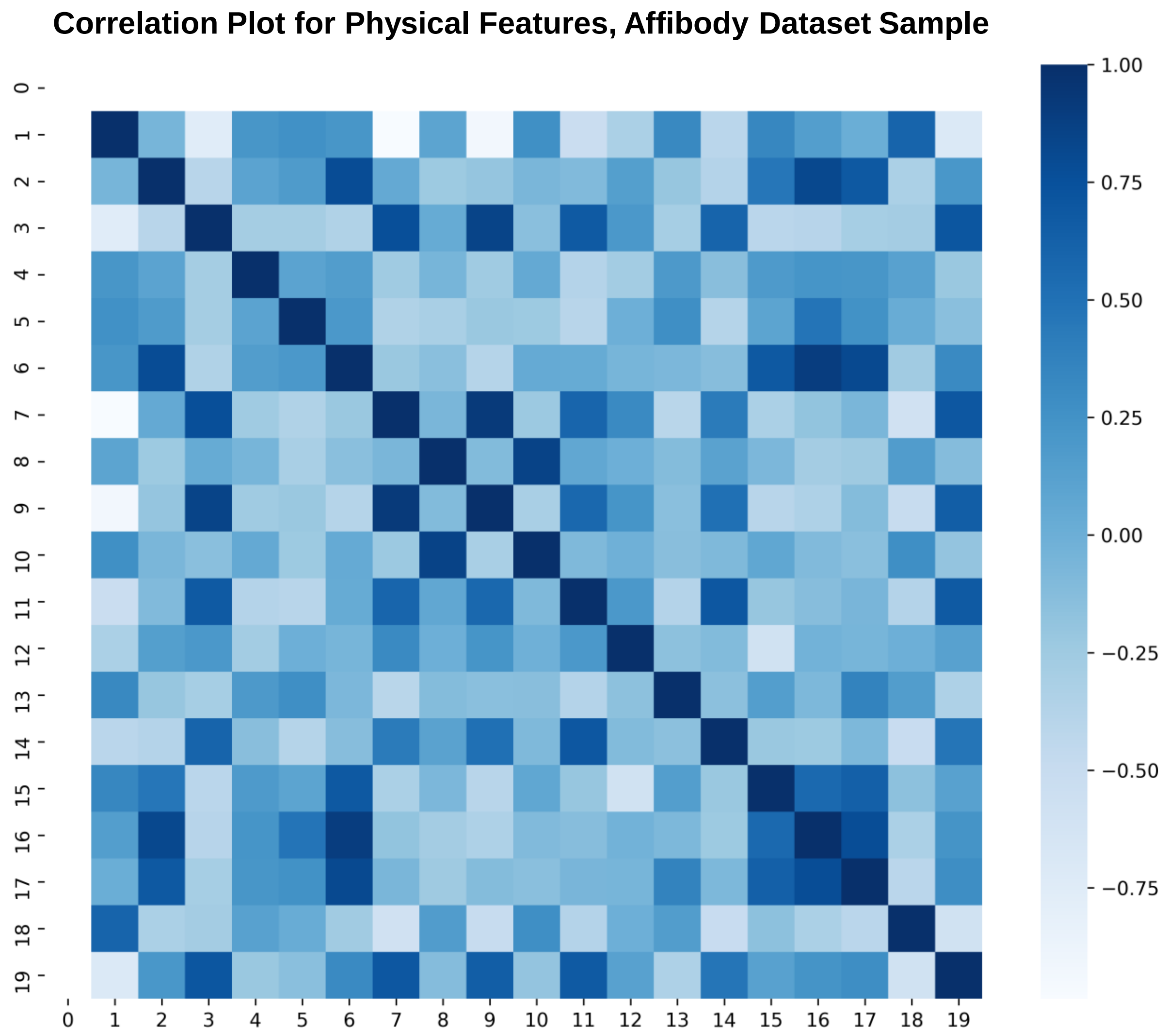


**Figure S1. Majority of physical features are at least correlated with one other physical feature; leaving us insights on attribute dependencies in the dataset.** The plot is drawn upon sampling from both naïve and enriched populations. Bold colors represent a higher correlation of features. As the affibody length in our dataset was fixed, the first row and column of the matrix are empty. Insights on the given correlation plot include i) a high correlation of Boman index and protein flexibility index, ii) a high correlation of aromaticity with molecular weight, MSW, and refractivity, iii) Aliphatic index having a correlation with multiple features such as bulkiness, H_Eisenberg, and H_Gravy.

**0)L, 1)Boman,2)Aromaticity, 3)Aliphatic,4)Instability, 5)Charge, 6)MW, 7)H_Eisenberg, 8)uH_Eisenberg, 9H)_GRAVY,10)uH_GRAVY, 11)Z3_1, 12)Z3_2, 13)Z3_3,14)Levitt alpha, 15)MSS, 16)MSW, 17)refractivity, 18)flexibility, 19)bulkiness**

#### Table S1: Figure 4 T-test Results

| | Comparison | P-Value |
|---|---|---|
| Under-sampling | Physical vs. **OH** | **2.37E-27** |
| | Physical vs. **UniRep** | **2.60E-24** |
| | Physical vs. **ESM** | **2.22E-20** |
| | OH vs. **UniRep** | **6.86E-06** |
| | **OH** vs. ESM | **9.31E-13** |
| | **UniRep** vs. ESM | **6.44E-08** |
| R-oversampling | Physical vs. **OH** | **6.12E-52** |
| | Physical vs**. UniRep** | **3.20E-50** |
| | Physical vs. **ESM** | **5.11E-50** |
| | OH vs. UniRep | 0.564 |
| | **OH** vs. ESM | **1.72E-18** |
| | **UniRep** vs. ESM | **2.19E-17** |
| SMOTE | Physical vs. **OH** | **2.06E-51** |
| | Physical vs. **UniRep** | **2.87E-50** |
| | Physical vs. **ESM** | **1.15E-50** |
| | OH vs. UniRep | 0.268 |
| | **OH** vs. ESM | **9.45E-16** |
| | **UniRep** vs. ESM | **2.76E-16** |

Table S2: Figure 5 T-test Results-part1

| Comparisons R-Oversampling for i vs. j | Mean i | Mean j | P-value |
|---|---|---|---|
| OH-vs-UniRep | 91.92 | 92 | 0.619375 |
| OH-vs-ESM | 91.92 | 89.91 | **1.06E-18** |
| OH-vs-UniRep+OH | 91.92 | 92.54 | **0.000201** |
| OH-vs-ESM+OH | 91.92 | 91.45 | 0.026281 |
| OH-vs-ESM+UniRep | 91.92 | 90.56 | **1.77E-09** |
| OH-vs-All | 91.92 | 92.61 | **8.38E-05** |
| OH-vs-Upvoted | 91.92 | 96.81 | **2.97E-31** |
| UniRep-vs-ESM | 92 | 89.91 | **2.04E-16** |
| UniRep-vs-UniRep+OH | 92 | 92.54 | 0.002124 |
| UniRep-vs-ESM+OH | 92 | 91.45 | 0.015463 |
| UniRep-vs-ESM+UniRep | 92 | 90.56 | **1.57E-09** |
| UniRep-vs-All | 92 | 92.61 | **0.000896** |
| UniRep-vs-Upvoted | 92 | 96.81 | **8.12E-26** |
| ESM-vs-UniRep+OH | 89.91 | 92.54 | **4.29E-19** |
| ESM-vs-ESM+OH | 89.91 | 91.45 | **2.15E-08** |
| ESM-vs-ESM+UniRep | 89.91 | 90.56 | **0.000344** |
| ESM-vs-All | 89.91 | 92.61 | **9.44E-19** |
| ESM-vs-Upvoted | 89.91 | 96.81 | **2.52E-38** |
| UniRep+OH-vs-ESM+OH | 92.54 | 91.45 | **1.47E-05** |
| UniRep+OH-vs-ESM+UniRep | 92.54 | 90.56 | **4.37E-13** |
| UniRep+OH-vs-All | 92.54 | 92.61 | 0.706828 |
| UniRep+OH-vs-Upvoted | 92.54 | 96.81 | **8.42E-24** |
| ESM+OH-vs-ESM+UniRep | 91.45 | 90.56 | **0.000355** |
| ESM+OH-vs-All | 91.45 | 92.61 | **7.00E-06** |
| ESM+OH-vs-Upvoted | 91.45 | 96.81 | **1.35E-19** |
| ESM+UniRep-vs-All | 90.56 | 92.61 | **2.59E-13** |
| ESM+UniRep-vs-Upvoted | 90.56 | 96.81 | **2.84E-25** |
| All-vs-Upvoted | 92.61 | 96.81 | **8.43E-23** |

Bonferroni Correction for Rejecting Null Hypothesis

$$\alpha = 0.05/28 \cong 0.002$$

Table S2: Figure 5 T-test Results-part 2

| **Comparisons for SMOTE for i vs. j** | **Mean i** | **Mean j** | **P-value** |
|---|---|---|---|
| OH-vs-UniRep | 92.88 | 93.07 | 0.222968 |
| OH-vs-ESM | 92.88 | 91.09 | **9.29E-15** |
| OH-vs-UniRep+OH | 92.88 | 92.53 | 0.050908 |
| OH-vs-ESM+OH | 92.88 | 91.61 | **5.80E-11** |
| OH-vs-ESM+UniRep | 92.88 | 90.47 | **1.85E-15** |
| OH-vs-All | 92.88 | 92.58 | 0.074292 |
| OH-vs-Upvoted | 92.88 | 97.08 | **4.02E-24** |
| UniRep-vs-ESM | 93.07 | 91.09 | **9.01E-17** |
| UniRep-vs-UniRep+OH | 93.07 | 92.53 | 0.003022 |
| UniRep-vs-ESM+OH | 93.07 | 91.61 | **3.23E-13** |
| UniRep-vs-ESM+UniRep | 93.07 | 90.47 | **1.72E-16** |
| UniRep-vs-All | 93.07 | 92.58 | 0.004057 |
| UniRep-vs-Upvoted | 93.07 | 97.08 | **6.46E-25** |
| ESM-vs-UniRep+OH | 91.09 | 92.53 | **2.43E-10** |
| ESM-vs-ESM+OH | 91.09 | 91.61 | **0.000153** |
| ESM-vs-ESM+UniRep | 91.09 | 90.47 | **0.000995** |
| ESM-vs-All | 91.09 | 92.58 | **1.50E-11** |
| ESM-vs-Upvoted | 91.09 | 97.08 | **1.88E-32** |
| UniRep+OH-vs-ESM+OH | 92.53 | 91.61 | **1.56E-06** |
| UniRep+OH-vs-ESM+UniRep | 92.53 | 90.47 | **8.55E-13** |
| UniRep+OH-vs-All | 92.53 | 92.58 | 0.792126 |
| UniRep+OH-vs-Upvoted | 92.53 | 97.08 | **1.11E-21** |
| ESM+OH-vs-ESM+UniRep | 91.61 | 90.47 | **1.11E-07** |
| ESM+OH-vs-All | 91.61 | 92.58 | **1.50E-07** |
| ESM+OH-vs-Upvoted | 91.61 | 97.08 | **2.07E-34** |
| ESM+UniRep-vs-All | 90.47 | 92.58 | **1.74E-13** |
| ESM+UniRep-vs-Upvoted | 90.47 | 97.08 | **1.37E-24** |
| All-vs-Upvoted | 92.58 | 97.08 | **4.06E-23** |

Bonferroni Correction for Rejecting Null Hypothesis

$$\alpha = 0.05/28 \cong 0.002$$

Table S3: SMOTE either Improved the performance or had no hampering effect with respect to R-Oversampling.

| Comparison for Samplings R-Oversampling vs. SMOTE | Mean R-Oversampling | Mean SMOTE | P-Value |
|---|---|---|---|
| OH | 91.92 | 92.88 | **7.30e-08** |
| UniRep | 92 | 93.07 | **3.08e-08** |
| ESM | 89.91 | 91.09 | **1.90e-11** |
| UniRep+OH | 92.54 | 92.53 | 0.96 |
| ESM+OH | 91.45 | 91.61 | 0.44 |
| ESM+UniRep | 90.56 | 90.47 | 0.65 |
| All | 92.61 | 92.58 | 0.88 |
| Upvoted | 96.81 | 97.08 | **0.002** |

Bonferroni Correction for Rejecting Null Hypothesis $\alpha = 0.05/8 \cong 0.006$

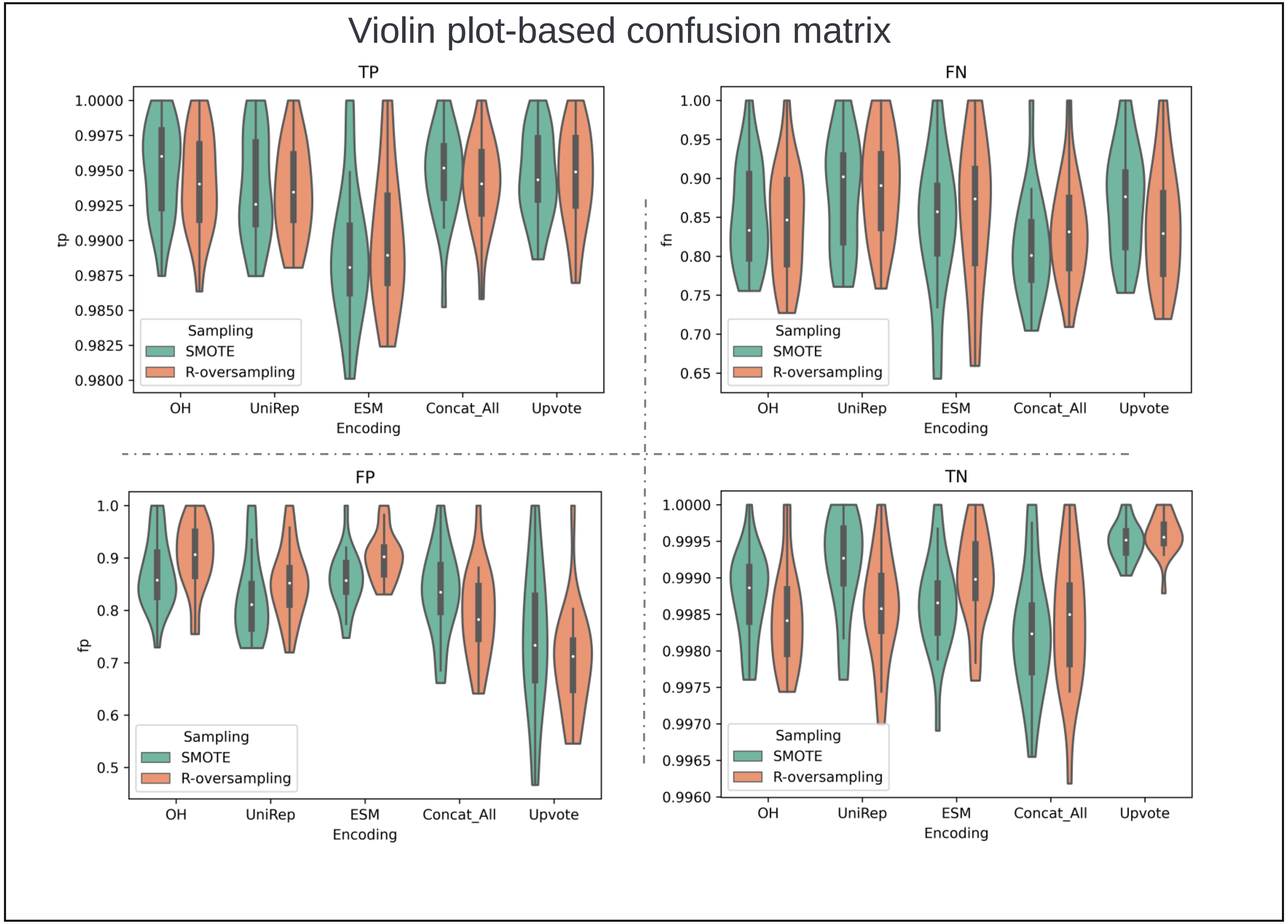


Figure S2. **Individual Encodings Perform uniquely in the confusion matrix entities.** While the overall predictive performance is the main goal and it is represented via F1-Score throughout the literature, inspecting how each model performs for maximizing true positives(TP) and true negatives(TN) while minimizing false positives(FP) and false negatives(FN), provides novel insights about each model performance.

### Table S4. T-Test for Figure S2

#### Stat for TP

| Encoding | P-value | Mean i | Mean j | Sampling |
|---|---|---|---|---|
| OH-vs-UniRep | 0.532 | 0.994 | 0.993 | R-Oversampling |
| OH-vs-ESM | **0.005** | 0.994 | 0.990 | R-Oversampling |
| OH-vs-Concat_All | 0.861 | 0.994 | 0.994 | R-Oversampling |
| OH-vs-Upvote | 0.848 | 0.994 | 0.994 | R-Oversampling |
| UniRep-vs-ESM | 0.016 | 0.993 | 0.990 | R-Oversampling |
| UniRep-vs-Concat_All | 0.634 | 0.993 | 0.994 | R-Oversampling |
| UniRep-vs-Upvote | 0.410 | 0.993 | 0.994 | R-Oversampling |
| ESM-vs-Concat_All | **0.006** | 0.990 | 0.994 | R-Oversampling |
| ESM-vs-Upvote | **0.003** | 0.990 | 0.994 | R-Oversampling |
| Concat_All-vs-Upvote | 0.707 | 0.994 | 0.994 | R-Oversampling |
| OH-vs-UniRep | 0.340 | 0.995 | 0.994 | SMOTE |
| OH-vs-ESM | **1.03E-4** | 0.995 | 0.989 | SMOTE |
| OH-vs-Concat_All | 0.764 | 0.995 | 0.995 | SMOTE |
| OH-vs-Upvote | 0.760 | 0.995 | 0.995 | SMOTE |
| UniRep-vs-ESM | **0.002** | 0.994 | 0.989 | SMOTE |
| UniRep-vs-Concat_All | 0.493 | 0.994 | 0.995 | SMOTE |
| UniRep-vs-Upvote | 0.480 | 0.994 | 0.995 | SMOTE |
| ESM-vs-Concat_All | **1.95E-4** | 0.989 | 0.995 | SMOTE |
| ESM-vs-Upvote | **1.65E-4** | 0.989 | 0.995 | SMOTE |
| Concat_All-vs-Upvote | 0.998 | 0.995 | 0.995 | SMOTE |

#### Stat for FN

| Encoding | P-value | Mean i | Mean j | Sampling |
|---|---|---|---|---|
| OH-vs-UniRep | 0.041 | 0.844 | 0.890 | R-Oversampling |
| OH-vs-ESM | 0.77 | 0.844 | 0.852 | R-Oversampling |
| OH-vs-Concat_All | 0.609 | 0.844 | 0.833 | R-Oversampling |
| OH-vs-Upvote | 0.88 | 0.844 | 0.840 | R-Oversampling |
| UniRep-vs-ESM | 0.152 | 0.890 | 0.852 | R-Oversampling |
| UniRep-vs-Concat_All | **0.01** | 0.890 | 0.833 | R-Oversampling |
| UniRep-vs-Upvote | 0.034 | 0.890 | 0.840 | R-Oversampling |
| ESM-vs-Concat_All | 0.47 | 0.852 | 0.833 | R-Oversampling |
| ESM-vs-Upvote | 0.679 | 0.852 | 0.840 | R-Oversampling |
| Concat_All-vs-Upvote | 0.734 | 0.833 | 0.840 | R-Oversampling |
| OH-vs-UniRep | 0.281 | 0.854 | 0.878 | SMOTE |
| OH-vs-ESM | 0.59 | 0.854 | 0.840 | SMOTE |
| OH-vs-Concat_All | 0.054 | 0.854 | 0.812 | SMOTE |
| OH-vs-Upvote | 0.463 | 0.854 | 0.870 | SMOTE |
| UniRep-vs-ESM | 0.153 | 0.878 | 0.840 | SMOTE |
| UniRep-vs-Concat_All | **0.005** | 0.878 | 0.812 | SMOTE |
| UniRep-vs-Upvote | 0.708 | 0.878 | 0.870 | SMOTE |
| ESM-vs-Concat_All | 0.265 | 0.840 | 0.812 | SMOTE |
| ESM-vs-Upvote | 0.251 | 0.840 | 0.870 | SMOTE |
| Concat_All-vs-Upvote | **0.01** | 0.812 | 0.870 | SMOTE |

Bonferroni Correction for Rejecting Null Hypothesis $\alpha = 0.05/5 \cong 0.01$

### Table S4. T-Test for Figure S2

#### Stat for TN

| Encoding | P-value | Mean i | Mean j | Sampling |
|---|---|---|---|---|
| OH-vs-UniRep | 0.569 | 0.9984 | 0.9986 | R-Oversampling |
| OH-vs-ESM | 0.018 | 0.9984 | 0.9990 | R-Oversampling |
| OH-vs-Concat_All | 0.758 | 0.9984 | 0.9984 | R-Oversampling |
| OH-vs-Upvote | **4.60E-07** | 0.9984 | 0.9996 | R-Oversampling |
| UniRep-vs-ESM | 0.081 | 0.9986 | 0.9990 | R-Oversampling |
| UniRep-vs-Concat_All | 0.429 | 0.9986 | 0.9984 | R-Oversampling |
| UniRep-vs-Upvote | **7.12E-6** | 0.9986 | 0.9996 | R-Oversampling |
| ESM-vs-Concat_All | 0.021 | 0.9990 | 0.9984 | R-Oversampling |
| ESM-vs-Upvote | **0.001** | 0.9990 | 0.9996 | R-Oversampling |
| Concat_All-vs-Upvote | **9.80E-06** | 0.9984 | 0.9996 | R-Oversampling |
| OH-vs-UniRep | 0.340 | 0.9987 | 0.9992 | SMOTE |
| OH-vs-ESM | **1.03E-4** | 0.9987 | 0.9986 | SMOTE |
| OH-vs-Concat_All | 0.764 | 0.9987 | 0.9982 | SMOTE |
| OH-vs-Upvote | 0.760 | 0.9987 | 0.9995 | SMOTE |
| UniRep-vs-ESM | **0.002** | 0.9992 | 0.9986 | SMOTE |
| UniRep-vs-Concat_All | 0.493 | 0.9992 | 0.9982 | SMOTE |
| UniRep-vs-Upvote | 0.480 | 0.9992 | 0.9995 | SMOTE |
| ESM-vs-Concat_All | **1.95E-4** | 0.9986 | 0.9982 | SMOTE |
| ESM-vs-Upvote | **1.65E-4** | 0.9986 | 0.9995 | SMOTE |
| Concat_All-vs-Upvote | 0.998 | 0.9982 | 0.9995 | SMOTE |

#### Stat for FP

| Encoding | P-value | Mean i | Mean j | Sampling |
|---|---|---|---|---|
| OH-vs-UniRep | 0.022 | 0.903 | 0.852 | R-Oversampling |
| OH-vs-ESM | 0.975 | 0.903 | 0.903 | R-Oversampling |
| OH-vs-Concat_All | **7.37E-05** | 0.903 | 0.794 | R-Oversampling |
| OH-vs-Upvote | **2.54E-08** | 0.903 | 0.703 | R-Oversampling |
| UniRep-vs-ESM | 0.01 | 0.852 | 0.903 | R-Oversampling |
| UniRep-vs-Concat_All | 0.025 | 0.852 | 0.794 | R-Oversampling |
| UniRep-vs-Upvote | **6.07E-06** | 0.852 | 0.703 | R-Oversampling |
| ESM-vs-Concat_All | **2.71E-05** | 0.903 | 0.794 | R-Oversampling |
| ESM-vs-Upvote | **2.10E-08** | 0.903 | 0.703 | R-Oversampling |
| Concat_All-vs-Upvote | 0.05 | 0.794 | 0.703 | R-Oversampling |
| OH-vs-UniRep | 0.501 | 0.872 | 0.824 | SMOTE |
| OH-vs-ESM | 0.131 | 0.872 | 0.858 | SMOTE |
| OH-vs-Concat_All | **0.001** | 0.872 | 0.834 | SMOTE |
| OH-vs-Upvote | 0.121 | 0.872 | 0.743 | SMOTE |
| UniRep-vs-ESM | 0.692 | 0.824 | 0.858 | SMOTE |
| UniRep-vs-Concat_All | 0.033 | 0.824 | 0.834 | SMOTE |
| UniRep-vs-Upvote | 0.291 | 0.824 | 0.743 | SMOTE |
| ESM-vs-Concat_All | **0.002** | 0.858 | 0.834 | SMOTE |
| ESM-vs-Upvote | 0.018 | 0.858 | 0.743 | SMOTE |
| Concat_All-vs-Upvote | 0.05 | 0.834 | 0.743 | SMOTE |

Bonferroni Correction for Rejecting Null Hypothesis $\alpha = 0.05/5 \cong 0.01$

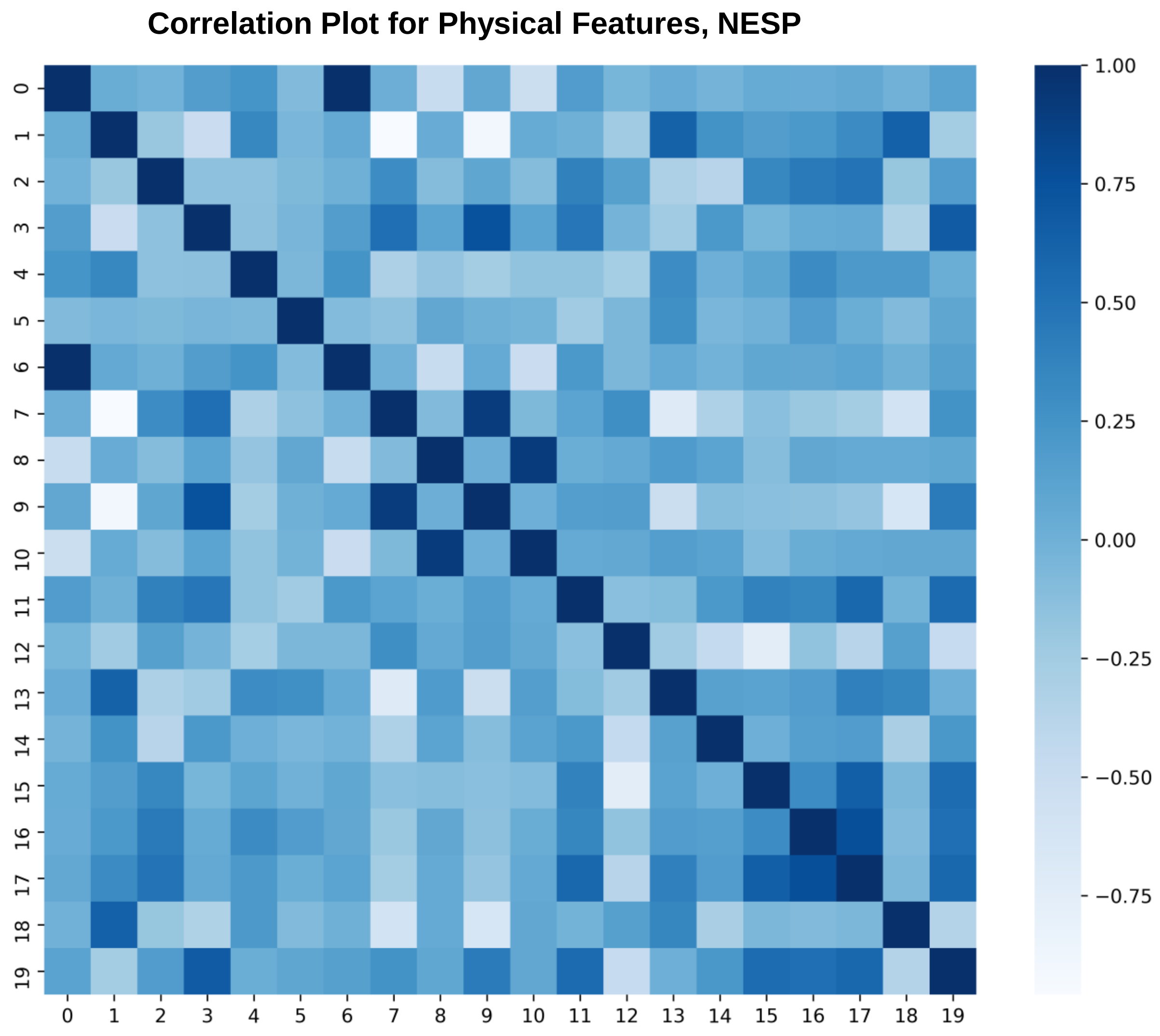


Note: From this Figure (Figure S2) the results are generated from NESP data**.**

**0)L, 1)Boman,2)Aromaticity, 3)Aliphatic,4)Instability, 5)Charge, 6)MW, 7)H_Eisenberg, 8)uH_Eisenberg, 9H)_GRAVY,10)uH_GRAVY, 11)Z3_1, 12)Z3_2, 13)Z3_3,14)levitt_alpha, 15)MSS, 16)MSW, 17)refractivity, 18)flexibility, 19)bulkiness**

Figure S3. Physical feature correlation plot for NESP dataset.

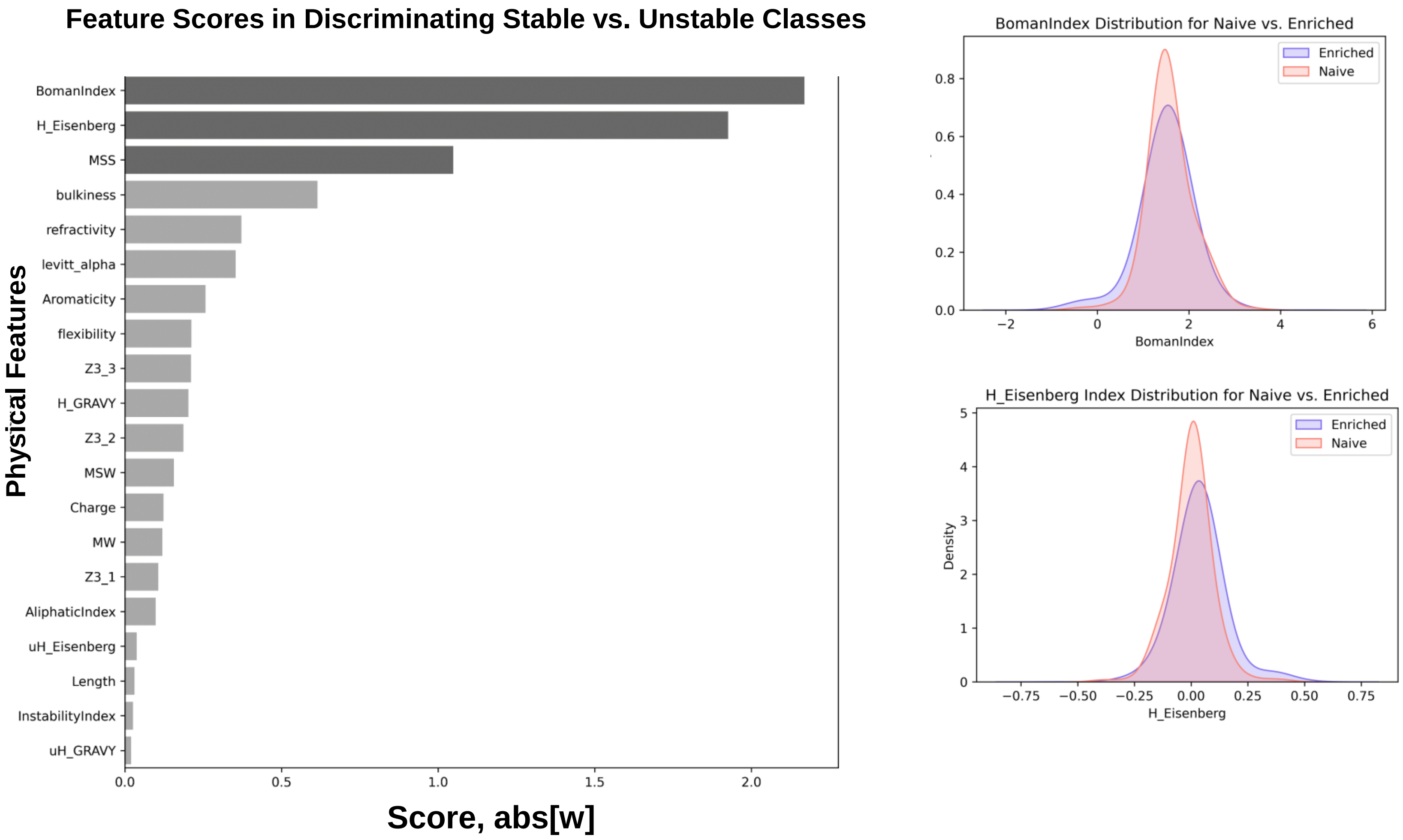


Figure S4: Physical feature ranking for NESP dataset. The feature ranking is presented after using all the data for the classification of stable vs. unstable sequences. Boman Index, H_Eisenberg, and MSS are the lead features. However, their scores are not significantly higher than the other physical attributes, indicating that more features incorporate in the final F1_Score=0.86.

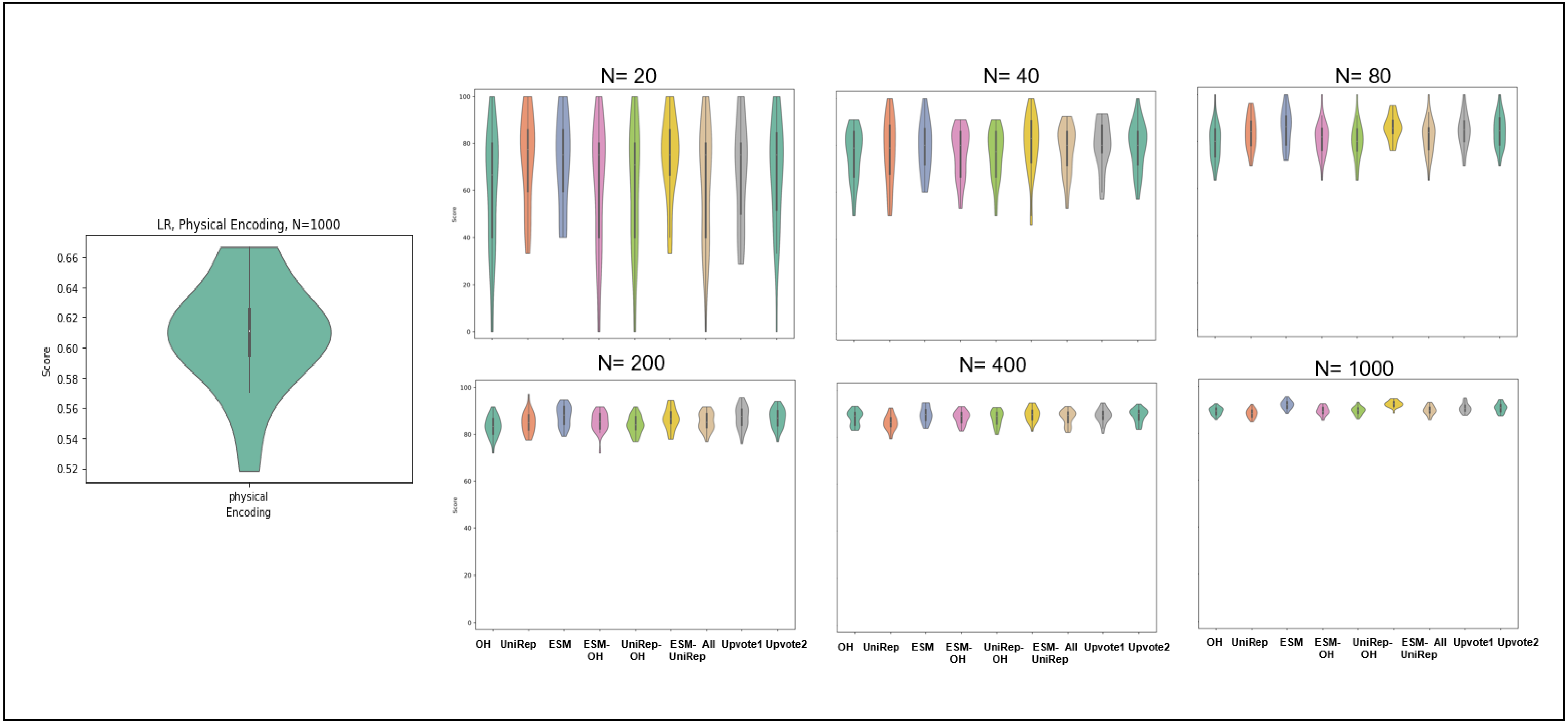


Figure S5. Physical Feature representation while using maximum N=1000, performed poorly and have not got selected for the main figure.

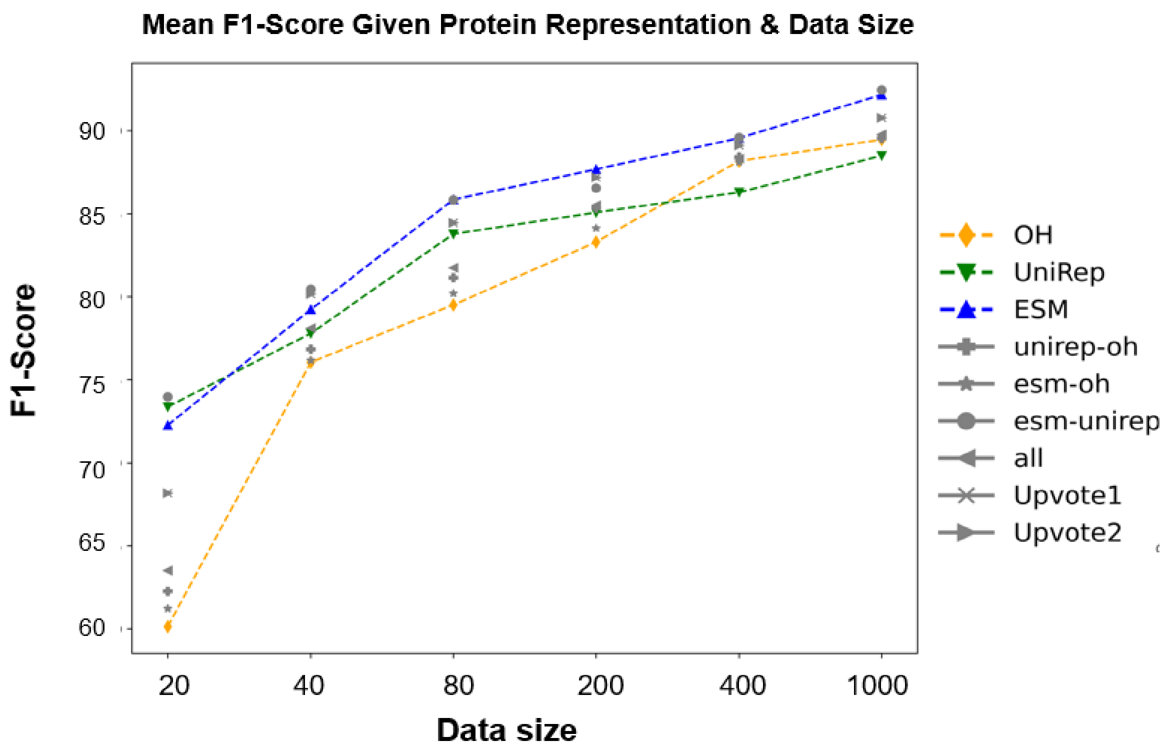


Figure S6. The predictive performances (F1-score) of multiple protein representations (One-Hot, UniRep, and ESM) were evaluated across stable (Tm>=60C; n=3140) vs. unstable (Tm<= 35C; n=1116) proteins. The performance of individual representations is compared against the effects of concatenating each embedding as well as ensemble methods (upvote1: hard voting, upvote2: soft voting). The violin plots were generated by repeating the analysis over 30 random seeds for sensitivity analysis. N represents the total number of data used with 0.3 as a test-size ratio. Welch t-test with unequal variances has been implemented over the obtained results to showcase the statistical significance in comparisons (refer to supplementary information for p-values). Highlight: In low data size (N=20), One-Hot performs poorly with the mean F1-score of 0.60, and the embeddings that included One-Hot were outperformed by both ESM and UniRep. While increasing the data size resulted in increased performance for all the methods, concatenating ESM with UniRep representations obtained the best score, with a mean F1 of 0.92. A complete statistical evaluation comparing each condition is provided in the supplementary information
